# Impaired neurogenesis and synaptogenesis in iPSC-derived Parkinson’s patient cortical neurons with D620N VPS35 mutation

**DOI:** 10.1101/2024.08.07.606995

**Authors:** Ute Scheller, ChoongKu Lee, Philip Seibler, Miso Mitkovski, JeongSeop Rhee, Christoph van Riesen

## Abstract

Presynaptic dysfunction is an important early process in the pathophysiology of Parkinson’s disease (PD) that drives disease progression. To gain insight into the intrinsic synapse impairment in PD, we performed comprehensive electrophysiological and morphological analysis of iPSC-derived cortical neurons derived from PD patients with the VPS35-D620N mutation. Our findings reveal significant impairment in neurogenesis and synaptogenesis within individual patient neurons, culminating to synaptic dysfunction even in the absence of neuronal interactions. The neurons exhibited significantly reduced synaptic responsiveness, fewer synapse and decreased dendritic length and complexity. Thus, the VPS35-D620N mutation in human cortical neurons independently can cause pathophysiology via synaptic dysfunction. Our study highlights the urgent need to develop disease-modifying therapies aimed at preserving synaptic function in PD.

## Introduction

Parkinson’s disease (PD) is the second most common neurodegenerative disease after Alzheimeŕs disease^1^. Recent cumulative evidence suggests that the pathophysiology of PD is initiated and driven by presynaptic mechanisms. This notion is supported by the fact that in post-mortem studies in human patients, 90% of aggregated alpha-synuclein is located presynaptically^2,3^. The neurodegenerative process is thought to start at the level of the synapse and then spread more proximally to the soma^4^. Imaging studies in early PD patients have shown a reduction in presynaptic markers, particularly in the substantia nigra pars compacta (SNpc) and dopamine transporter in the striatum suggesting that presynaptic pathology may precede dopaminergic cell death^5,6^. Aberrant cortico-striatal activity and degeneration of cortico-striatal glutamatergic synapses have been proposed as independent stressors for dopaminergic neurons in the SN, suggesting at least some disease propagation from the cortex downwards^7,8^. Cortico-striatal and nigrostriatal terminals form a highly interactive complex with medium spiny neurons (MSNs) in the striatum. It has been argued that the focal symptom onset of PD reflects the somatotopic organization of the cortex and the striatum and is therefore better explained by an early cortical input-driven pathology than by random dopaminergic degeneration in the non-somatotopically organized SNpc^7^. In addition, transcranial magnetic stimulation (TMS) has demonstrated that alterations in cortical plasticity and cortical disinhibition occur at an early stage in PD, representing either a compensatory or a maladaptive change that contributes to disease progression^9–11^. Dopaminergic neurons have been described as particularly vulnerable due to cell-autonomous factors, including their high arborization with long axons, high firing rate with broad spikes and slow Ca^2+^-oscillations among others^12^, which may contribute to the neurodegeneration in complex PD pathophysiological changes. Dysfunction of the cortical input onto the basal ganglia system may be an additional stressor leading to dopaminergic degeneration in PD.

Despite the recent insights into the importance of presynaptic mechanisms in the pathophysiology of PD, there questions remain many unanswered. The lack of animal models for sporadic PD has slowed progress in this field. Consequently, animal models carrying monogenic mutations that lead to PD in humans are of significant value. Monogenic mutations have been identified in 10% of PD patients^1^, including a rare pathogenic autosomal dominant variant D620N in the vacuolar sorting protein 35 (VPS35)^13,14^. Patients with VPS35 mutations develop late-onset levodopa-responsive parkinsonism that is clinically indistinguishable from sporadic PD^15^. The precise mechanism by which VPS35 mutations, including the D620N mutation, cause degeneration of the nigrostriatal dopamine and other neurotransmitter systems in humans remains elusive^16^, largely due to a paucity of comprehensive data on their impact on neuronal function. VPS35 is part of the retromer complex, which is involved in the endosomal transport system^16,17^. The VPS35-D620N mutation has been described to affect endosomal-lysosomal pathways^18^, impair mitochondrial function^19^, and to be associated with synaptic dysfunction^16^. VPS35 is part of the retromer complex, which is involved in the endosomal transport system^16,17^. The VPS35-D620N mutation has been described to affect endosomal-lysosomal pathways^18^, impair mitochondrial function^19^, and to be associated with synaptic dysfunction^16^.

This study aimed to further elucidate the role of VPS35 in human PD, with a particular focus on synaptic mechanisms. We used an autaptic neuronal culture system to enable the detailed study of the intrinsic electrophysiological and morphological properties of iPSC-derived VPS35 D620N cortical single neurons from PD patients. In the absence of any external stimuli, we observed that neurogenesis (dendritic complexity and length), synaptogenesis and synaptic strength were impaired in the patient-derived cell lines. These unique properties of each mutant neuron have the potential to significantly impact the neuronal network in PD, thereby highlighting their importance in causing synaptic pathology.

## Methods

### Cell lines

The iPSC lines have been characterized previously and were derived from dermal fibroblasts of two healthy donors (SFC086-03-01, C1; SFC084-03-02, C2) and two Parkinson’s disease patients (iPS-L6731-7, PD1; iPS-L6634-5, PD2) carrying a heterozygous c.1858G>A (p.D620N) missense mutation in the *VPS35* gene (Supplementary Table S1). The cell lines are abbreviated from here on as C1, C2, PD1 and PD2, respectively. After the completion of the experiments in our lab, routine karyotypic analysis of the iPSC cultures in another laboratory revealed chromosomal abnormalities in the C2 control line after long-term passaging (SFC084-03-02: 46,XX,del(10)(p12p12)[15]/45,idem,-X[5]). As our experiments had already been completed and no cells were left, we were unable to determine whether the C2 cells used carried the karyotypic change.

All participants provided written informed consent (Ethics Committee that approved this part of the study: University of Lübeck, Lübeck, Germany). All iPSC lines were cultured on Matrigel (BD Bioscience) coated dishes in mTeSR1 medium (STEMCELL Technologies).

### Differentiation into cortical neurons

The direct differentiation of iPSCs into cortical neurons was performed as previously described ^20,21^. Briefly, iPSCs were dissociated into single cells using Accutase (Life Technologies) and replated onto Matrigel-coated dishes. Differentiation was initiated by adding the SMAD pathway inhibitors dorsomorphine (Tocris; 1μM) and SB 431542 (Tocris; 10μM) to the cultures. From day 13 to 17, the cells were cultured in neural maintenance medium (NMM)^20^ containing basic fibroblast growth factor (R&D Systems; 20ng/ml) and brain-derived neurotrophic factor (BDNF, STEMCELL Technologies; 20ng/ml). Neural rosettes were replated on day 18 in NMM with BDNF, glial cell-derived neurotrophic factor (GDNF, STEMCELL Technologies; 20ng/ml), and ascorbic acid (Sigma-Aldrich; 0.2mM). The medium was changed every other day until day 27. On day 28, the rosettes were dissociated using Accutase and plated in NMM supplemented with BDNF, GDNF, and ascorbic acid. The medium was replaced every 2-3 days until day 43. On day 43, BDNF, GDNF, and ascorbic acid were removed, and the cells were cultured in NMM until day 44//49.

### Cell Culture

Human iPSC (hiPSC) were seeded onto prepared mouse astrocyte feeder cells according to previously established protocols^22^. Briefly, mouse astrocytes were obtained from dissected cortex of P0 wild-type animals, enzymatically digested with trypsin-EDTA and cultured in DMEM (GIBCO) for 7-10 days. 6-well culture plates with coverslips were prepared to allow the growth of astrocyte micro-islands. Glass coverslips were coated with agarose and stamped with a customized stamp to generate islands coated with poly-D-lysine, acetic acid and collagen^23^. hIPS were thawed in a water bath at 37°C, diluted in an iPS medium containing ROCK inhibitor (48% DMEM/F12, 48% Neurobasal medium, 1% N2, 1% B27, 0.1% beta-mercaptoethanol, 0.5% L-glutamine, 0.5% NEAA, 0.5% FBS, 0.1% ROCK inhibitor), filtered through a 40µm cell strainer (BD Falcon) and counted using tryptan blue staining. Appropriate numbers of cells were seeded onto the prepared astrocyte microislands and incubated at 37°C for 24 hours. The complete iPS-medium was replaced with fresh medium without ROCK inhibitor the next day and changed once per week for the following culture period. hiPSCs were seeded onto prepared mouse astrocyte feeder cells according to previously established protocols^22^. Briefly, mouse astrocytes were obtained from dissected cortex of P0 wild-type animals, enzymatically digested with trypsin-EDTA and cultured in DMEM (GIBCO) for 7-10 days. 6-well culture plates with coverslips were prepared to allow the growth of astrocyte micro-islands. Glass coverslips were coated with agarose and stamped with a customized stamp to generate islands coated with poly-D-lysine, acetic acid and collagen^23^. hIPS were thawed in a water bath at 37°C, diluted in an iPS medium containing ROCK inhibitor (48% DMEM/F12, 48% Neurobasal medium, 1% N2, 1% B27, 0.1% beta-mercaptoethanol, 0.5% L-glutamine, 0.5% NEAA, 0.5% FBS, 0.1% ROCK inhibitor), filtered through a 40µm cell strainer (BD Falcon) and counted using tryptan blue staining. Appropriate numbers of cells were seeded onto the prepared astrocyte microislands and incubated at 37°C for 24 hours. The complete iPS-medium was replaced with fresh medium without ROCK inhibitor the next day and changed once a week for the following culture period.

### Electrophysiology

Autaptic neurons were whole-cell patch clamped either using either a Multiclamp 700B (Axon Instruments, Molecular Devices) or an EPSC10 (HEKA electronics) amplifier controlled by Clampex software (Axon Instruments, Molecular Devices) or Patchmaster 2 (HEKA electronics), respectively. The extracellular bath solution (140mM NaCl, 4mM CaCl2, 4mM KCl, 10mM HEPES, 24mM MgCl2, 10mM glucose, adjusted to pH 7.3, and ∼310msOsmol/l) was maintained at a constant flow and pharmacological solutions were added as required using a custom-made flow pipette. Glass pipettes were made using a Sutter 2000 filament-based horizontal puller and their resistance ranged between 2.5 and 3.5 MΩ. The patch pipette solution consisted of 136 mM KCl, 17.8 mM HEPES, 1 mM EGTA, 4.6 mM MgCl2, 4 mM NaATP, 0.3 mM Na2GTP, 15 mM creatine phosphate, and 5 U/ml phosphocreatine kinase (315-320 mOsmol/l), pH 7.4. Neurons were visualized using an inverted microscope (Zeiss or Olympus). Only neurons with series resistance <15MΩ were analysed. For voltage clamp recordings cells were held at −70mV. In voltage mode synaptic responses and voltage-gated channel currents were evoked by electrical stimuli with a frequency of 0.2Hz and a stepwise (10mV) increase in membrane holding potentials from −80 to +70mV for 500ms each. To block glutamatergic and GABAergic receptors 10µM NBXQ (HelloBio) and 10µM bicuculline (Tocris) were added to the extracellular solution using a flow pipette aimed at the cell and the combined voltage-gated sodium and potassium channel current (I_total_) was measured. To isolate I_K_, 300nM TTX (Tocris) was applied exogenously and then 1mM 4-AP was added to record I_DR_. I_Na_ was calculated by subtracting I_K_ from I_total_ and A-type I_K_ (I_A_) was calculated by subtracting I_DR_ from I_K_. The densities of all currents were normalized to cell capacitance (pF). Using the current clamp mode R_input_ was calculated from membrane potential chance by current injection from −100pA to 0pA in 20pA intervals for 1s each. Action potentials (AP) were evoked by stepwise increasing (+10pA) current injection starting from 0pA. The patch pipette solution used to record these membrane properties consisted in 138mM K-gluconate, 16.8mM HEPES, 10mM NaCl, 1 mM MgCl_2_ 6H2O, 0.25mM EGTA, 4mM ATP-Mg^2+^, 0.3mM GTP–Na^+^ (solution adjusted to pH 7.4, 320mOsmol/litre). All experiments were performed at room temperature. All recordings were analyzed using AxographX software (version 1.5.4., Axograph Scientific).

### Immunofluorescence staining and imaging

Human autaptic neurons were washed with PBS, fixed with 4%PFA for 15 min, washed with PBS, quenched with 50mM glycine for 10 min, washed again with PBS and incubated with iT-FX image enhancer (Life Technologies) for 30 min. Cells were then permeabilized with 0.1% Triton X-100/2.5% NGS/PBS for 40 min, blocked with 2.5% NGS/PBS and incubated with primary antibodies for 1 hour. The primary antibodies used were: MAP2 (chicken, 1:1000, Novus Biological), SHANK2 (guinea pig, 1:250, Synaptic Systems), Synapsin 1/2 (rabbit, 1:500, Synaptic Systems). Cells were washed with 0.1% Triton X-100/2.5% NGS/PBS and incubated with secondary antibodies for 45 min. The following secondary antibodies were used: Alexa Flour 405 (anti-chicken 1:1000, Abcam), Alexa Flour 488 (anti-guinea pig 1:1000, Thermo Fisher), Alexa Flour 568 (anti-rabbit 1:1000, Thermo Fisher). After a final wash with 0.1% Triton X-100/2.5% NGS/PBS and PBS, cells were mounted with Aqua-Poly Mount (Polyscience). All steps were carried at room temperature. Fluorescence images were captured using a Leica TCS SP8 X confocal laser scanning microscope (Leica Microsystems).

### Image processing

Overlapping z-stacks (x,y,z: 229, 229, 299 nm) were acquired with the inverted Leica TCS SP8 X confocal laser scanning microscope equipped with a white light laser, motorized stage and a 63x 1.4 NA oil immersion objective. The Napari v. 0.4.18^24^ Noise2Void plugin^25^ was used to train a denoising model which was then applied to the corresponding channels of the original confocal images. For quality control purposes, the denoised channels were then merged with the original channels, reconstructed and calibrated using a custom written FIJI^26^ macro and then converted using the Imaris File Converter (v. 10.1.0). The Imaris image visualization and analysis software package (v. 10.1.0) was used for image analysis. Synapses were counted by colocalization between pre- and postsynaptic puncta labelled by Synapsin 1/2 and Shank2 respectively. Puncta were also required to colocalize with MAP2 dendritic staining. The Imaris plugin Filament-tracer was used in a semi-automatic mode to detect the dendritic tree marked by MAP2 staining and to calculate the total dendrite length (sum), the number of dendritic branching points and Sholl intersections (Sholl analysis).

### Statistical analysis

Statistical analysis and visualization were performed using GraphPad Prism (version 10.2.2, GraphPad Software). Data were checked for normality and, in the case of non-normal distribution, logarithmized (ln) prior to further analysis. To account for the factors ‘group’ (control vs PD), ‘subgroup’ (C1, C2, PD1, PD2) and ‘time’ (week 2-6), 3-way ANOVA was used to calculate differences between groups ‘control’ and ‘PD’. Additional subgroup comparisons were made using 2-way ANOVA and supplemented by Tukey’s test for multiple comparisons. There was no significant interaction between ‘subject’ and ‘time’ for any of the parameters analyzed. Group data are presented as mean ± SEM or median (IQR) for non-parametric data respectively. For visualization purposes, group comparisons (PD vs control) are presented as box plots (Tukey), where the means (or medians respectively) of the time points for each subgroup were calculated first, followed by the medians of the time points per group. Similarly, for subject comparisons, means or medians of time points were calculated first and presented as box plots (Tukey). All subgroup comparisons are shown in Supplementary Figures and Supplementary Table 3. Statistical results presented in box plots were calculated using 3-way or 2-way ANOVA. The significance level was set at p<0.05 (*≤0.05, **≤0.01, ***≤0.001, ****≤0.0001).

## Results

In this study, two neuronal cell lines derived from two PD patients carrying the VPS35 D620N mutation were compared with two cell lines from controls. Direct differentiation of iPSCs into cortical glutamatergic neurons was performed as previously described^20,21^. For the analysis, the human neurons were seeded into an autaptic culture system, in which single neurons were placed on individual mouse astrocyte feeder islands. The autaptic culture system allows reproducible characterization of single hiPSC derived neurons, as we and others have recently demonstrated^22,27^. Our analysis included electrophysiological patch-clamp recordings and morphological analysis by immunocytochemistry imaging over a period of 6 weeks.

### Neuronal differentiation and growth: active and passive membrane properties

First, we used whole-cell voltage and current patch-clamp recordings to measure passive and active membrane properties to verify proper neuronal development. Neuronal differentiation of hiPSCs could be demonstrated by typical morphological and membrane properties from 2 weeks in vitro (WIV) counted starting from the day of cultivation in the autaptic culture system.

### Passive membrane properties

Measurements of passive membrane properties (capacitance, resting membrane potential (RMP) and input resistance (R_input_)) were used as parameters for cell development. Capacitance, as a measure of cell size, remained stable over 6 WIV (Fig 1a, Supplementary Table 2) and there was no statistical difference between PD and control neurons (Fig. 1b). RMP and R_input_ as markers for cell excitability decreased slightly but not significantly over time indicating cell maturation (Fig. 1c, e, Supplementary Table 2). Control cell lines had a significantly lower RMP than PD patient cell lines (*p*=0.0119, Fig. 1d, Supplementary Table 2) with C1 being hyperpolarized (subgroup analysis, Supplementary Table 3, Supplementary Fig. 1b). R_input_ showed no significant difference between the VPS35-D620N patient and control cells (Fig. 1f, Supplementary Table 2).

**Figure 1.**
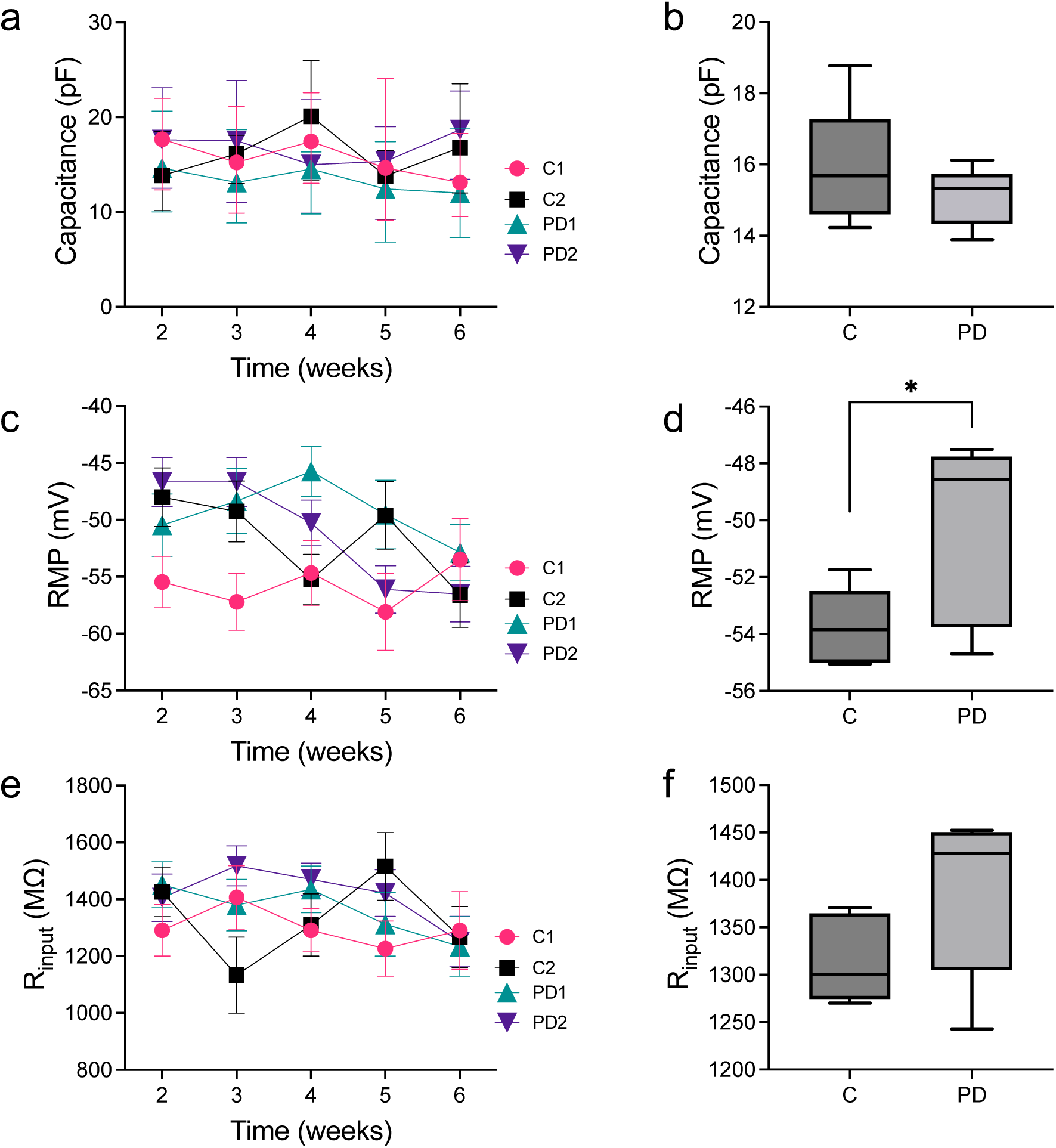
Passive Membrane Properties. **(a)** Cell capacitance for each subgroup and timepoint over 6 WID. Medians (IQR) are shown. (**b)** Group comparison of cell capacitance between control and PD cell-lines. (**c)** RMP for each subgroup over time. (**d**) Group comparison reveals controls had significantly lower RMP in control cell-lines. (**e)** Rinput per subgroup over time **d** and **f** group comparison between control vs PD. In graphs **c** and **e** means ± SEM are shown.

### Active membrane properties: Voltage-gated sodium and potassium channels

Next, we characterized active membrane properties by analyzing ion channel activity and action potential (AP) generation. First, we assessed the activity and density of voltage-gated sodium channels. The peak I_Na_ densities of PD and control neurons were identical (Fig. 2c, Supplementary Table 2). Similarly, the densities of I_K_, delayed rectifier K^+^-currents (I_DR_) and fast transient A-type K^+^-currents (I_A_) were not different between PD and control groups (Fig. 2f, i, l, Supplementary Table 2). Although I_Na_ and I_A_ densities were statistically different over time (‘time’ factor: *p*=0.0428, *p*=0.0028, Supplementary Table 2), there was no clear developmental pattern (Fig. 2b, e, h, k).

**Figure 2.**
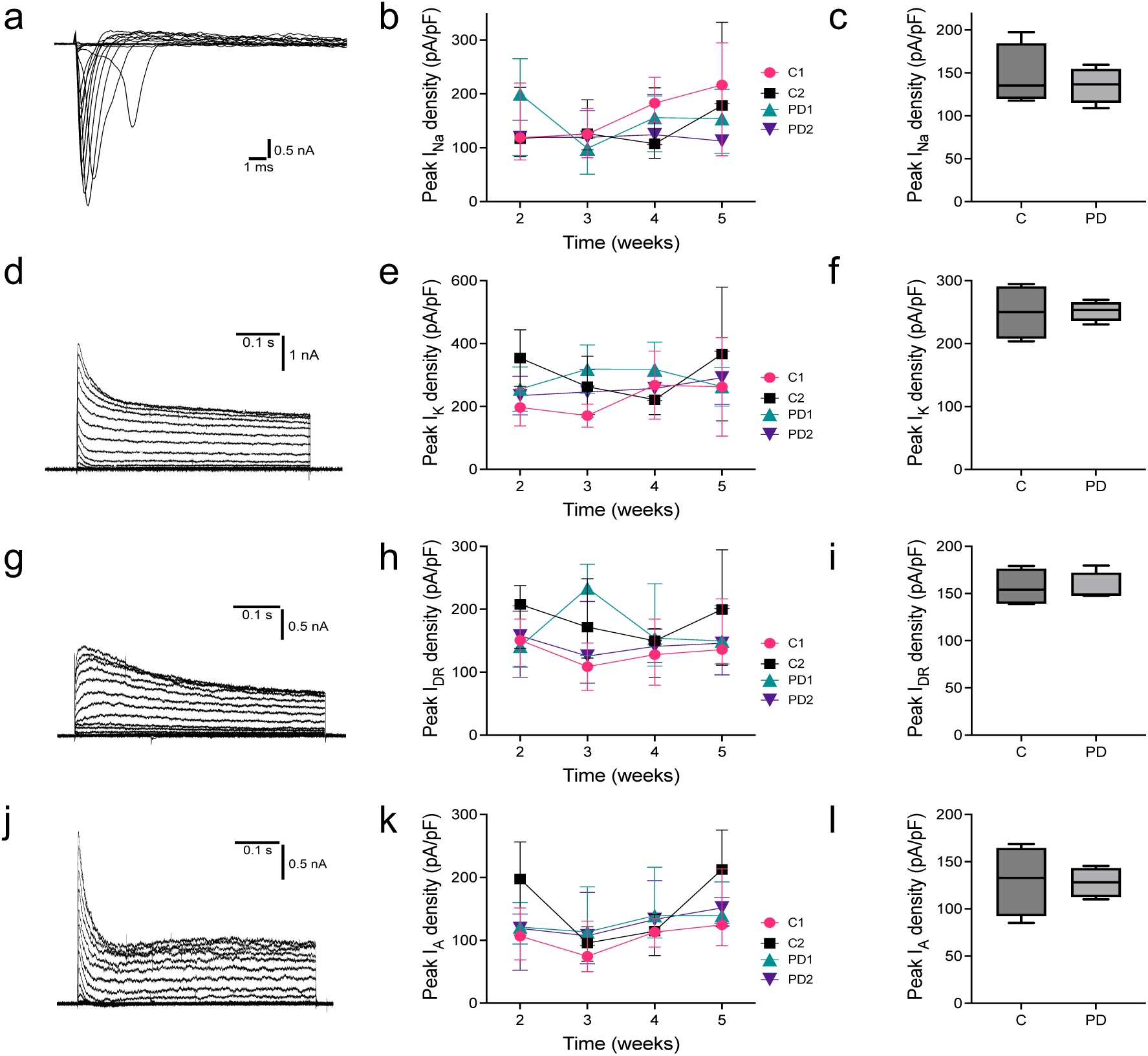
Voltage-gated Channels. Voltage-gated sodium and potassium currents are shown as peak current densities**. (a**, **d**, **g** and **j)** Representative traces showing depolarization-induced I_Na_, I_K_, I_DR_ and I_A_, respectively. (**b**, **e**, **h**, and **k)** Peak current densities over time for each subgroup. For all graphs medians (IQR) are given. (**c**, **f**, **i**, and **l)** Group comparison of peak current densities control vs PD cell-lines.

### Active membrane properties: action potentials

The hiPSC-derived neurons, which contained properly functioning voltage-gated ion channels, were found to generate AP. We characterized the APs by amplitude, duration, half-width, threshold, overshoot, afterhyperpolarisation (AHP), rheobase and firing pattern. The hiPSC-derived neurons were only analyzed, if normal ion channel function could be demonstrated. Six different AP firing patterns could be distinguished: abortive, immature, spontaneous, tonic, phasic I and phasic II (Supplementary Fig. 3a). To compare the numbers of APs, we selected the highest number from each cell recording. At the group level, PD neurons generated a significantly higher number of APs compared to controls (Fig. 3b). This effect was likely due to a decrease in the number of APs at WIV 5 and 6, predominantly in controls, which was reflected in a higher proportion of phasic I and absent tonic firing cells (Fig. 3a, Supplementary Fig. 3b, c, Supplementary Table 2). Control C1 also showed more phasic II and less spontaneous and tonic firing overall which might have further increased the difference (Supplementary Fig. 3b, Supplementary Table 2). No significant differences were found between PD and control cell lines for rheobase, AP threshold, AP overshoot, AP amplitude, AP duration and AP half-width (Fig. 3d, f, h, j, l, n, Supplementary Table 2). The AHP amplitude was significantly smaller in PD neurons (*p*=0.0279, Fig. 3p, Supplementary Table 2). The different AP parameters remained stable over time in both groups (Fig. 3c, i, k, m, o), except for the AP threshold and AP overshoot which decreased slightly over time (‘time’ factor: *p*<0.0001, *p*=0.0044, Fig. 3e, g, Supplementary Table 2). AP duration and half-width decreased minimally but not significantly indicating further AP maturation (Fig. 3k, m).

**Figure 3.**
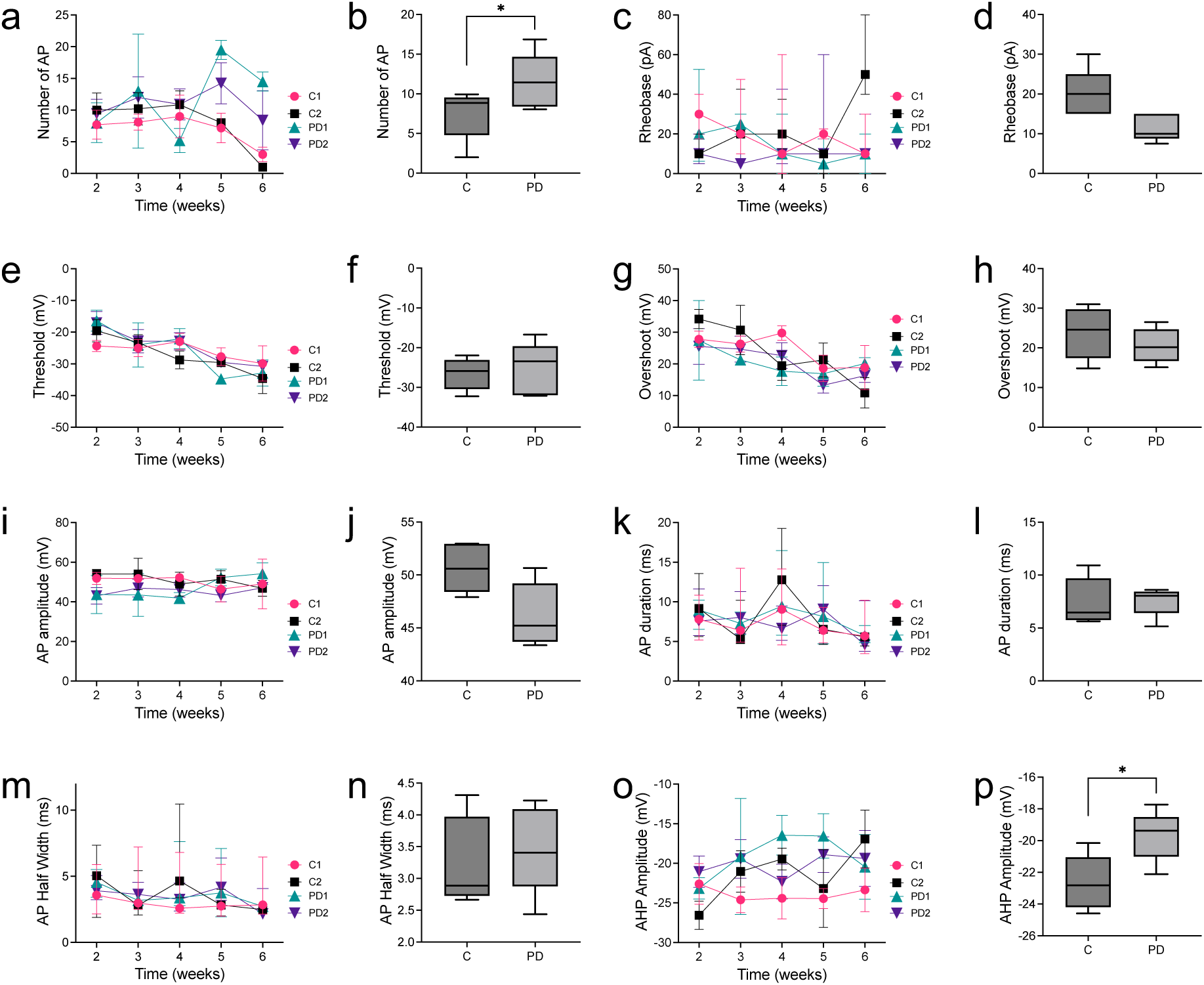
Active Membrane Properties. Various parameters generating and shaping action potential (AP) are shown along with the iPSC-derived neuron development (**a, c, e, g, I, k, m, o**) and in group comparison (**b, d, f, h, j, l, n, p**). Peak AP-number (**a** and **b**), Rheobase (**c** and **d**), Threshold (**e** and **f**), Overshoot (**g** and **h**), AP amplitude (**i** and **j**), AP duration (**k** and **l**), AP half width (**m** and **n**) and AHP amplitude (**o** and **p**) over WID. In **a**, **k** and **m** median (IQR), in **c, e, g, i** and **o** means ±SEM are depicted.

Overall, the neurons matured by 2 WIV with little further growth thereafter. Active and passive membrane properties indicated proper neuronal differentiation in all cell lines with only minor differences between PD and control cells.

### Impairment of synaptic release in PD neurons

We recorded evoked postsynaptic currents (eEPSC) to assess the development of functional synapses. The patient hiPSCs were significantly less likely to develop synaptic responses than those derived from controls (*p*<0.0001, Fig. 4c, Supplementary Table 2). In both groups the percentage of cells with a synaptic response increased over time (‘time’ factor: *p*=0.0043, Fig. 4b, Supplementary Table 2). While control neurons reached a steady state at 4 WIV, PD neurons showed a much slower increase. At 6 WIV, 65% and 69% of control cells developed a synaptic response whereas only 50% and 43% of PD cells did so. Further analysis of the subgroups (C1, C2, PD1, PD2) showed that the synaptic response was lower in each PD subject compared to each control (Supplementary Fig. 5a, Supplementary Table 3). In addition, EPSC amplitudes were significantly lower in PD neurons (*p*<0.0001, Fig. 4f, Supplementary Table 2). There was no significant change in EPSC amplitude over time (Fig. 4e). Each control subject had higher EPSC amplitudes than PD patient cells (Supplementary Fig. 5b, Supplementary Table 3). Thus, PD neurons were less likely to form functional synapses and had smaller eEPSC amplitudes than controls.

**Figure 4:**
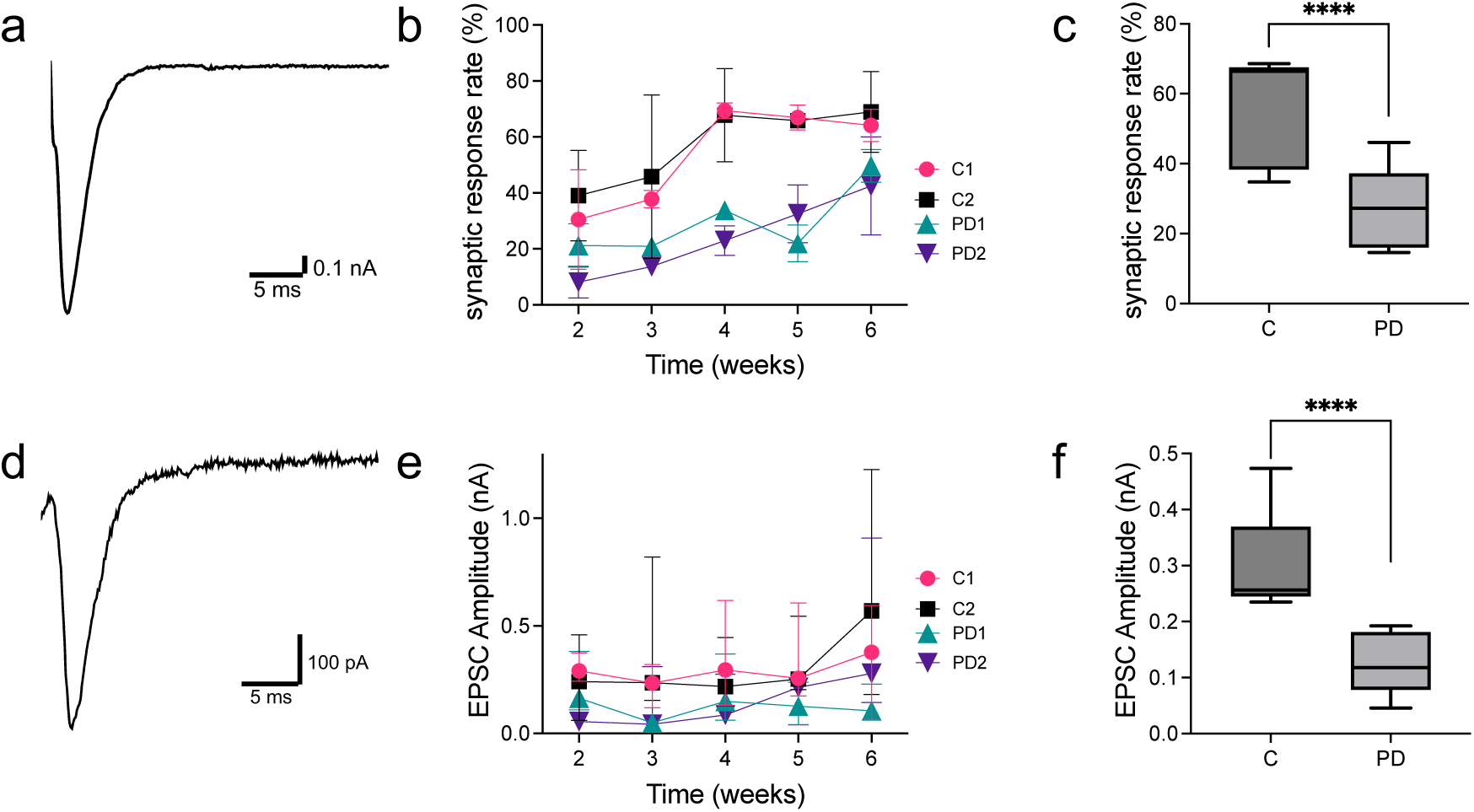
Synaptic Response. (**a**, **d)** Representative traces of EPSC from control (**a**) and PD (**d**) neurons at 4 WID each. (**b** and **e**) The proportion of neurons synaptic release and EPSC amplitude over time. (**c** and **f**) Group comparisons for the proportion of neurons with synaptic release and EPSC amplitude control versus PD patient cell lines. In **e** median (IQR) and in **b** means ±SEM are given.

### Morphological analysis: filament tracing and synapse counting

We used immunocytochemistry to monitor neurogenesis and synaptogenesis. Dendritic length and complexity were analyzed by MAP2 staining (filament tracing, Sholl analysis). In general, cells showed some diversity in morphological shape, with no marked differences between groups and time points (Fig. 5c). When comparing the two groups, it was found that control neurons had a greater total summed dendrite length than PD cells (*p*=0.0014, Fig. 5b, Supplementary Table 2). To varying degrees, the dendrite length tended to increase over time in all groups up to 5 weeks (‘time’ factor, *p*=0.0013, Fig. 5a, Supplementary Table. 2). To examine the dendritic complexity, we analyzed the number of dendritic branching points and Sholl intersections. While the number of dendritic branching points did not differ between groups (Fig. 5e, Supplementary Table 2), the number of Sholl intersections was significantly reduced in PD neurons compared to controls, indicating a higher complexity of the control cell lines (*p*=0.0006, Fig. 5h, Supplementary Table 2). The number of dendritic branching points increased by 3 weeks and fluctuated by 6 WIV, while the dendritic length increased up to 5 WIV (‘time’ factor *p*=0.0105, *p*=0.0043, Fig. 5d, g, Supplementary Table. 2).

**Figure 5.**
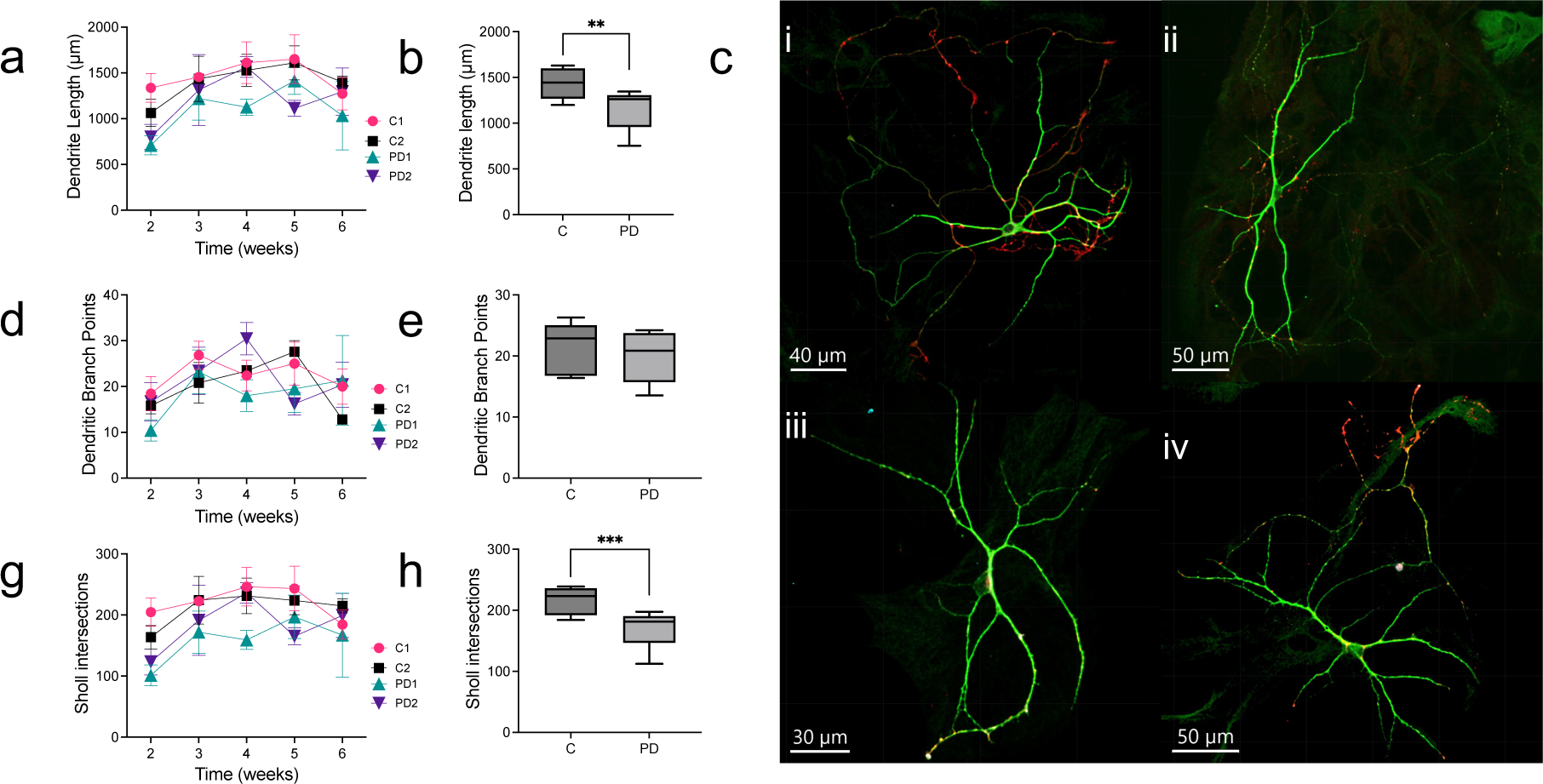
Morphological Analysis. (**a**, **d**, **g**) Dendritic filament length (sum), number of branching points and average numbers of dendrites intersecting Sholl circles up to 200 μm distance per subgroup over time. (**b, e, h**) Group comparison between control and PD patient cell-lines. (**c**) Representative fluorescence images for human autaptic neurons for each subgroup at 4 WID. **ci** and **cii** are control neurons and **ciii** and **civ** are neurons induced from PD cell-lines. For immunohistochemical staining, MAP2 (green) was used as dendritic, Synapsin1/2 (red) as presynaptic and Shank2 (blue) as postsynaptic marker.

Next, we wanted to investigate whether the reduced synaptic response rate was due to reduced synaptogenesis in general or due to a reduced number of functional synapses. We analyzed colocalized pre- and postsynaptic puncta labelled with the presynaptic marker Synapsin 1/2 and the postsynaptic marker Shank2, to identify them as synapses (Fig. 6c, d). The PD groups showed a significant decrease in the total number of synapses compared to the control neurons (*p*<0.0001, Fig. 6b, Supplementary Table 2). In addition, there were significant differences in number of synapses during neuronal development. In the control neurons, the number of synapses increased significantly up to WIV 4 and then decreased slightly at WIV 6 (‘time’ factor *p*=0.0003, Fig. 6a, Supplementary Table 2). In the PD groups, however, there were no obvious changes in synapse numbers from week to week. There was a clear dissociation between the rate of neuronal development and the rate of new synapse formation (‘time’ factor *p*=0.0003, Fig. 6a, Supplementary Table 2). Subgroup comparisons confirmed significant differences between all control and PD cell lines (Supplementary Fig. 5c, Supplementary Table 3).

**Figure 6.**
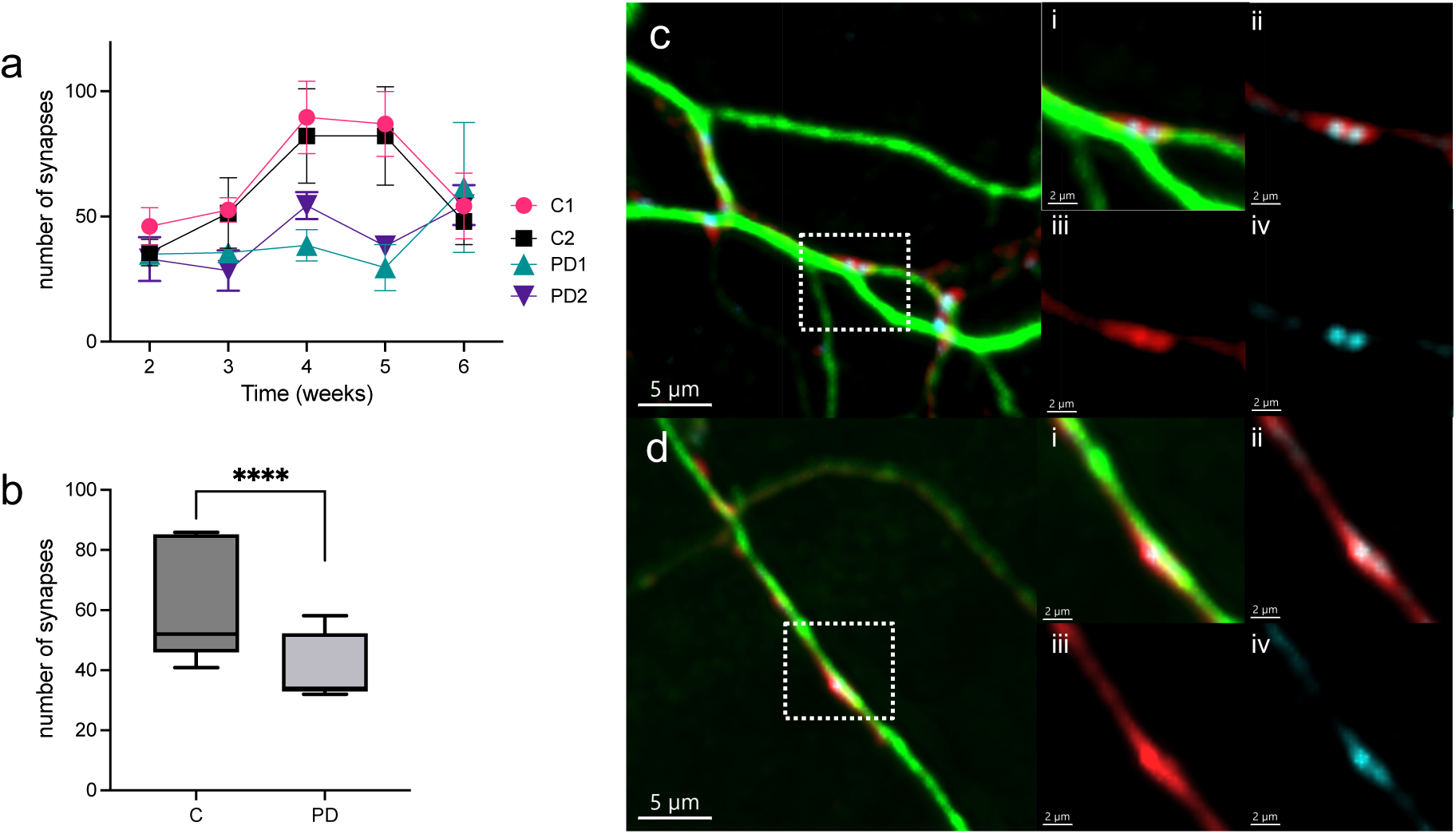
Number of synapses. **(a)** Development of the number of synapses per subgroup over time. (**b**) Group comparison of controls vs PD patient cell-lines. (**c, d**) Representative fluorescence images for human autaptic neurons for each subgroup at 4 WID. Synapse number were counted by colocalizing Synapsin1/2 puncta (presynaptic marker, red) and Shank2 puncta (postsynaptic marker, blue) within proximity to dendrites (MAP2, green). Control cell (**c**) and PD patient cell (**d**) with magnification of synapse: **i** MAP2, Synapsin 1/2, Shank2, **ii** Synapsin 1/2, Shank2; **iii** Synapsin 1/2; **iv** Shank2.

In conclusion, our study demonstrated significant impairments in neurogenesis, marked by reduced dendritic length and complexity, and in synaptogenesis, indicated by a decreased number of synapses and functional synapses, in hiPSC-derived neurons from PD patients with the D620N VPS35 mutation compared to controls.

## Discussion

Modelling an age-related human neurodegenerative disease such as PD presents numerous challenges. To address this, we utilized human cortical iPSC-derived neurons from two PD patients carrying the VPS35 D620N mutation as a simplified cellular model and monitored them for six weeks. To date, only a few electrophysiological studies of human, predominantly dopaminergic iPSC-derived neurons from PD patients, have been conducted using large-scale culture systems. However, these culture systems present a challenge in distinguishing whether observed phenotypes are due to the neurons themselves or from interactions with neighboring neurons. The autaptic culture system excludes inter-neuronal communication, thus allowing us to focus exclusively on the intrinsic properties of the targeted neurons. As a result, the phenotypes observed in VPS35 D620N hiPSC-derived neurons were not influenced by extraneous factors such as dopaminergic stimulation or environmental fluctuations, rendering them invaluable for elucidating the aberrant neuronal network characteristic of Parkinson’s patients.

### Growth pattern

The control cell line C2 developed a karyotypic change (46,XX,del(10)(p12p12)[15]/45,idem,-X[5]) after long-term passaging. Although there is some heterogeneity between the control groups in terms of membrane properties and morphological shape, we did not observe any significant difference between them in general. Furthermore, heterogeneity in cellular iPSC-derived models is a well-known issue, which emphasizes the importance of using multiple, or in the best case, isogenic controls^28^.

### Impaired neurogenesis

Given that VPS35 plays a key role within the retromer complex, which is critical for endosomal protein sorting and intracellular transport^16,17^, it is plausible to hypothesize that the VPS35-D620N mutation potentially contributes to Parkinson’s disease through various disruptive cellular mechanisms. The mutation has been associated with impaired neurogenesis^29–31^. The published phenotypic manifestations of the D620N VPS35 mutation in the literature so far such as dopaminergic cell death^29,30^, reduced neuronal proliferation, neurite degeneration, impaired neurite outgrowth and dendritic complexity^31^, show some variability and remain inconclusive. For example, one study investigating the effect of the PD-associated VPS35 mutation on dopaminergic hiPSC-derived neurons revealed reduced differentiation efficiency and increased apoptosis compared to controls^29^, whereas in another study human dopaminergic neurons derived from PD patients with the VPS35 mutation differentiated into dopaminergic neurons similar to controls^32^. Inconsistencies have also been observed in VPS35-D620N knock-in mouse models (VKI). While Chen et al. observed degeneration of dopaminergic neurons and neurites in certain brain regions (animals aged 13 months)^30^, others found no such degeneration or changes in younger animals (aged 3 months) or in primary cortical cell cultures from VKI mice (DIV 21)^33,34^. This variability may be influenced by experimental models and conditions, and the timing of assessments, amongst others. Highlighting the importance of long-term observations to detect age-related phenotypes consistent with the delayed dopaminergic degeneration observed in PD patients, a recent study found dopaminergic cell loss and elevated dopamine levels only in older animals (aged 15/16 months) but not before^35^.

Beyond the dopaminergic system, a recent study examined neurogenesis in the hippocampal neural progenitor pool of transgenic mice carrying the VPS35-D620N mutation^31^. The study unveiled a diminished pool of neural progenitors, alongside reduced proliferation and migration, coupled with impaired neurite outgrowth characterized by diminished length and complexity in hippocampal neurons, signifying impaired neurogenesis^31^. Similar morphological changes have been observed in neurons derived from PD patients with other genetic mutations, including diminished neurite length and complexity in IPSC-derived cells harboring Parkin and LRRK2 mutations, as well as reduced soma size and neuronal extensions in cells from idiopathic PD patients^36–40^.

In our study, we found that human cortical iPSC-derived neurons with the VPS35-D620N mutation also showed impaired neurogenesis, characterized by reduced dendritic length and cell complexity.

These findings suggest that impaired neurogenesis may represent a general PD phenotype, that extends beyond dopaminergic cells. Further investigation is essential to unravel the contributions of these pathways to neurodegeneration and to elucidate the neurogenesis deficits seen in cellular models of PD.

### Impaired synaptogenesis and reduced number of functional synapses

The presence of VPS35 at synapses underscores its significance in synaptic function^41,42^. Numerous studies have explored synaptic dysfunction in VPS35-deficient models, with a particular focus on cortical and dopaminergic neurons of animal models. This research has identified changes in synapse quantity^41,43^, spine formation^43^, synaptic vesicle dynamics^43,44^, neurotransmitter release^33,45^ and clustering and surface expression of AMPA receptors^41,43^, indicating a broad association of VPS35 with various synaptic functions. However, the findings relating to synaptic morphology and vesicle number have been inconsistent, potentially due to differences in experimental models (animal, KD, VKI), conditions (large-scale culture), and timing of assessments^33,34,42^. At the single cell level, our findings revealed significantly lower proportions of neurons with a synaptic response, reduced EPSC amplitudes and fewer synapses in IPSC-derived cortical neurons from PD patients carrying the VPS35 D620N mutation, suggesting impaired synaptogenesis in human VPS36 D620N cell lines.

Previous studies have not demonstrated a clear correlation between changes in synapse numbers and mEPSC size^34,41,43^. In our investigation, spanning from WIV 2 to 6, we observed that while synapse numbers in control neurons increased steadily in tandem with neuronal development, the EPSC size remained relatively stable, with only a slight increase at WIV 6. This indicates that despite an increase in synaptic formation during neuronal maturation, the absolute number of functional synapses remained largely unchanged. However, after 6 weeks, a reduction in synapse numbers paradoxically corresponded with an increase in the absolute number of functional synapses, leading to an enhanced EPSC size. It is likely that the period between WIV 2 and 6 is critical for functional synaptic conversion during neuronal development, though such conversion also occurs even when synapse numbers were lower. In D620N VPS35 mutants, the number of synapses fluctuated slightly over time but remained relatively stable with a small increase at week 6. Correspondingly, EPSC amplitudes showed fewer fluctuations than controls. Overall, PD patient-derived neurons showed a decrease in the total number of synapses and a reduction in neurons with synaptic responses compared to control, highlighting a significant impairment in synaptic function.

Therefore, we speculate that the abnormalities in synaptogenesis caused by D620N VPS35 mutants not only disrupt synapse formation but also selectively impair the conversion of structural synapses into functional ones. Consequently, PD-derived neurons exhibit not only a reduced number of synapses but also a diminished rate of functional synapse conversion.

The underlying molecular mechanisms of these synaptic anomalies likely involve disruptions in endosomal trafficking^46^. In particularly, VPS35 has been associated with early and late endosomes^47^, which are essential for synaptic vesicle recycling^48^ and AMPA receptor trafficking^49^. In human dopaminergic iPSC-derived neurons carrying the VPS35 D620N mutation, VPS35 co-localised with Rab5 and Rab7, markers of early and late endosomes, and showed reduced velocity and lower fission- and fusion-frequency of Rab5- and Rab7-vesicles compared to controls^29^. Thus, impaired trafficking of endocytosed SV could lead to an abnormal vesicle pool and altered expression of neurotransmitter receptors on plasma membranes, resulting in synaptic dysfunction. Our data in D620N mutant neurons suggest that VPS35 influences the formation of individual synapses and is differentially involved in the establishment of functional synapses in each neuron. However, the precise mechanisms by which the VPS35 mutation affects synaptogenesis, resulting in reduced synapse numbers and synaptic dysfunction, remains to be fully elucidated.

Moreover, synaptic dysfunction has also been implicated in other PD genes, including SNCA, LRRK2, Parkin and PINK1 amongst others^8,50–53^. For example, significantly reduced synaptic vesicle density was found in IPSC-derived dopaminergic neurons from a PD patient with a LRRK2 mutation^54^. However, data on synaptic transmission in IPSC derived PD neurons are scarce and show incongruent changes of sEPCS^55,56^. Furthermore, genome-wide studies in PD have identified differentially expressed genes associated with mitochondrial functions and protein degradation but also with synaptic vesicle dynamics and synaptic transmission^57,58^. This broader involvement of synaptic dysfunction underscores its pivotal role in the pathophysiology of PD, potentially even preceding dopaminergic neurodegeneration, as suggested by early imaging studies^5,6^.

## Conclusion

In conclusion, we have identified, for the first time, impaired neurogenesis and synaptogenesis in human glutamatergic iPSC-derived neurons from PD patients with the VPS35-D620N mutation. This discovery establishes a definitive synaptic phenotype in cortical neurons in a human-derived PD model. To uncover the precise mechanisms behind impaired neurite outgrowth and synaptic dysfunction, further research should investigate the synaptic vesicle pool, synaptic plasticity, and postsynaptic receptor density in greater detail. The establishment of robust in vitro PD models is hampered by the complex nature of the basal ganglia, a critical structure in PD pathophysiology, comprising various interacting cell types, nuclei, input- and output structures. Although our single-cell model is by default oversimplified, our results support the emerging notion of early presynaptic dysfunction beyond just dopaminergic neurons in PD. Enhancing our understanding of PD pathomechanisms and developing valid models for pharmacological testing are essential for enabling potential new treatments. Continued research is crucial to deepen our understanding of PD pathomechanisms and explore possible interventions.

## Acknowledgment

We thank Anja Günther for her excellent technical support during experiments and Albrecht Sigler for his expertise and excellent support during imaging analysis. US was funded by a scholarship from the Deutsche Forschungsgemeinschaft (DFG, German Research Foundation) - 413501650.

## Authors contribution

JSR, CvR, US conceived study and designed experiments. PS generated and provided human iPSC-derived neurons. US, CKL performed experiments. US, CKL, MM analysed data. JSR, CvR and US wrote the paper. All authors were involved final editing of the paper.

## Supplementary Figure Legends

**Supplementary Figure 1.**
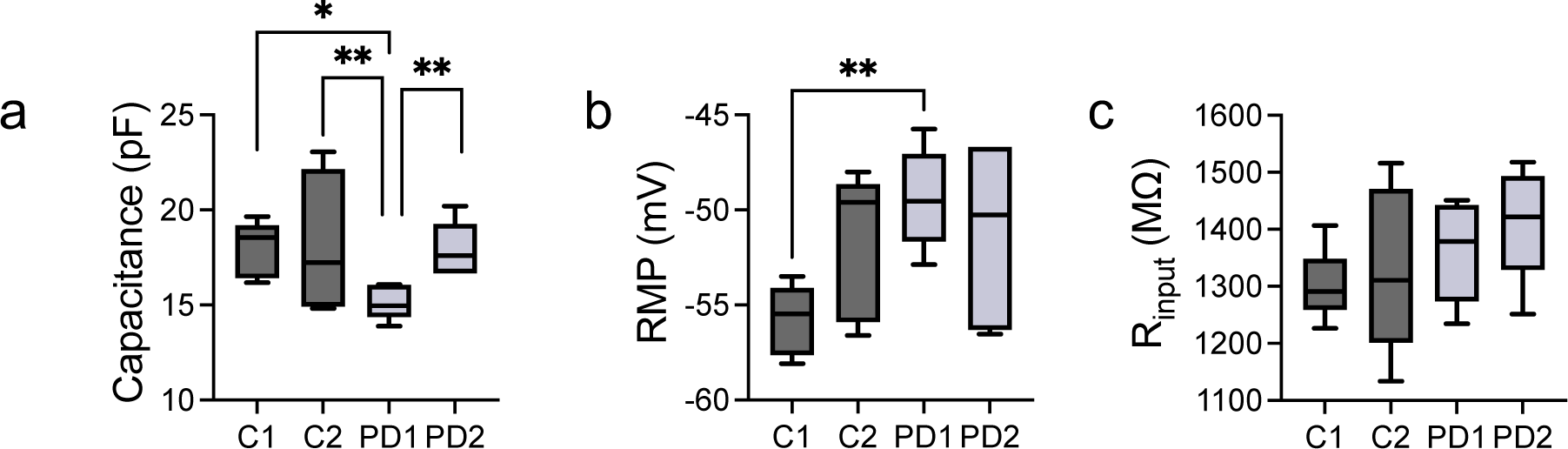
Passive Membrane Properties. Graphs show subgroup comparison (C1, C2, PD1, PD2) of passive membrane properties. (**a**) Capacitance; ‘subgroup’ factor: *p* =0.0041, C1 vs. PD1: *p* =0.0465, C2 vs PD1: *p* =0.0096, PD1 vs PD2: *p* =0.0084, 2-way ANOVA with multiple comparison test Tukey). (**b**) RMP; ‘subgroup’ factor: *p* = 0.0128, C1 vs PD1: *p* = 0.0069, 2-way ANOVA with multiple comparison test Tukey. (**c)** R_input_; there are no significant differences between subgroups.

**Supplementary Figure 2.**
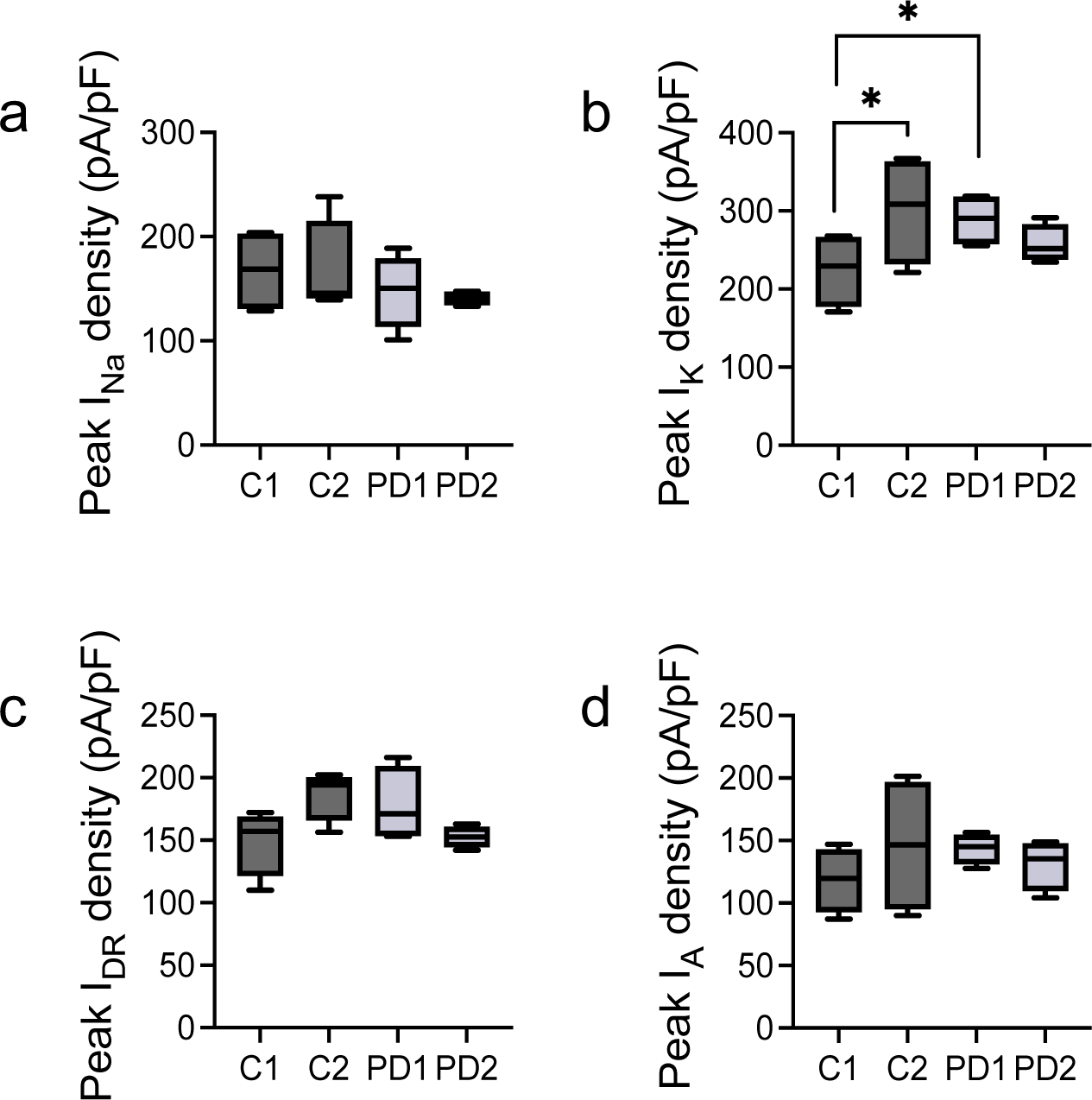
Voltage-gated Channels. Voltage-dependent Sodium and Potassium currents presented as peak densities per subgroups. **(a)** Graph for I_Na_ densities per subgroup; (**b-d**) Graph for total I_K_ (b), I_DR_ and I_A_ densities; Slight subgroup differences can be seen for peak I_K_ currents (’subgroup’ factor: P= 0.0094, C1 vs C2: *p* = 0.019, C1 vs PD1: *p* = 0.0182, 2-way ANOVA with multiple comparison test Tukey).

**Supplementary Figure 3.**
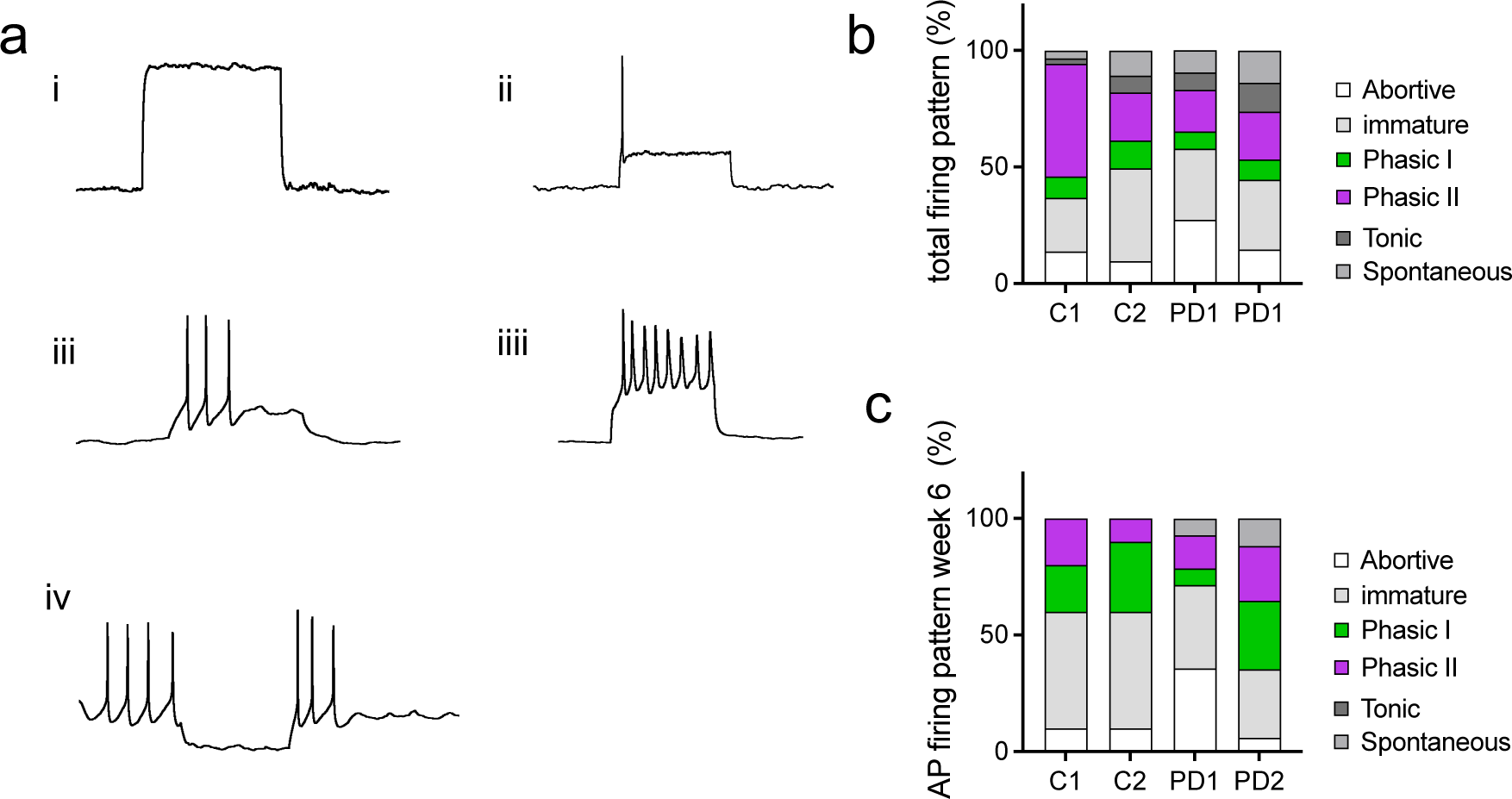
Action potential patterns. **(a)** Identified patterns of AP firing: abortive (**i**), phasic I (**ii**), phasic II (**iii**), tonic (**iiii**) and spontaneous (**iv**). Combined percentage of AP firing pattern per subgroup (**b**) and AP firing pattern of each subgroup at 6 WID (**c**). At 6 WID, no tonic and in controls no spontaneous firing cells were recorded.

**Supplementary Figure 4.**
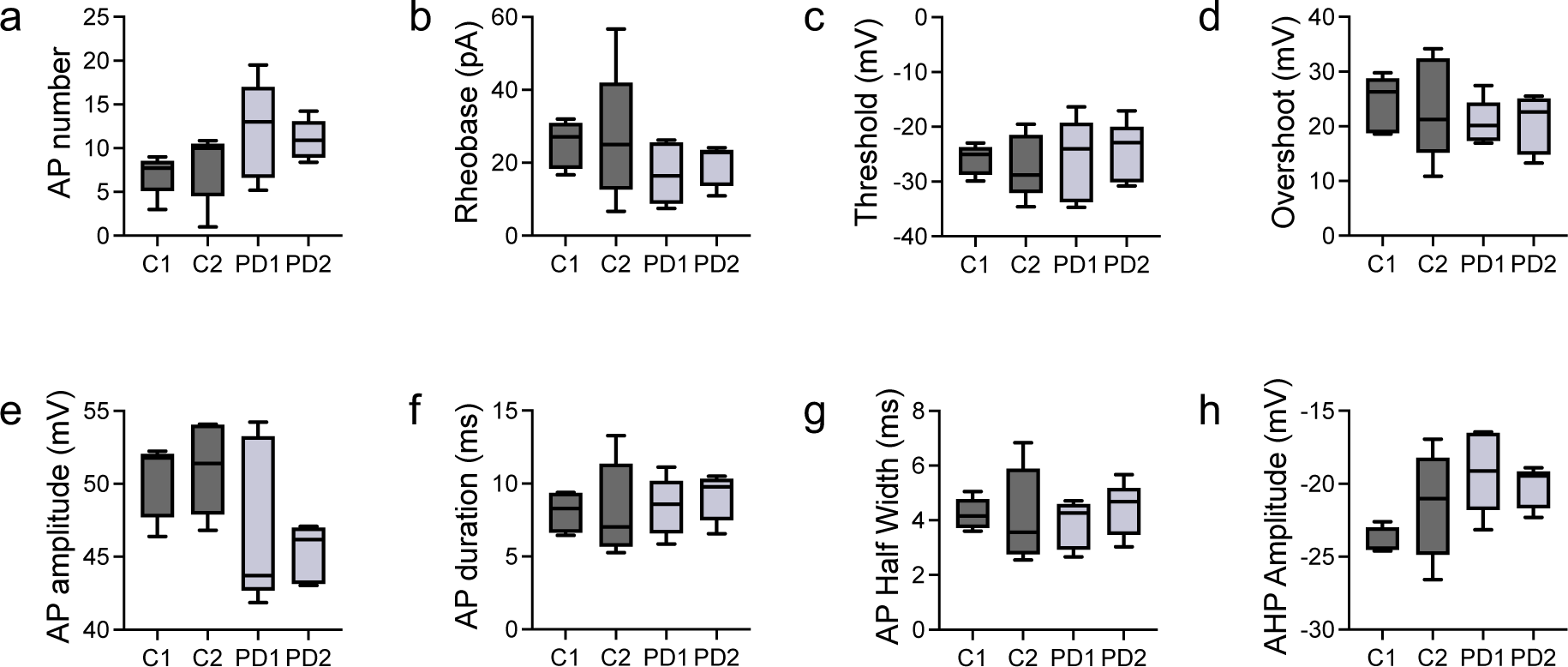
Active Membrane Properties; Parameters for action potential. **(a-d)** Subgroup comparison for various parameters of action potential shape. AP numbers (**a**) and AP-amplitudes (**e**) are slightly lower, respectively higher, in controls vs PD patient cell-lines.

**Supplementary Figure 5.**
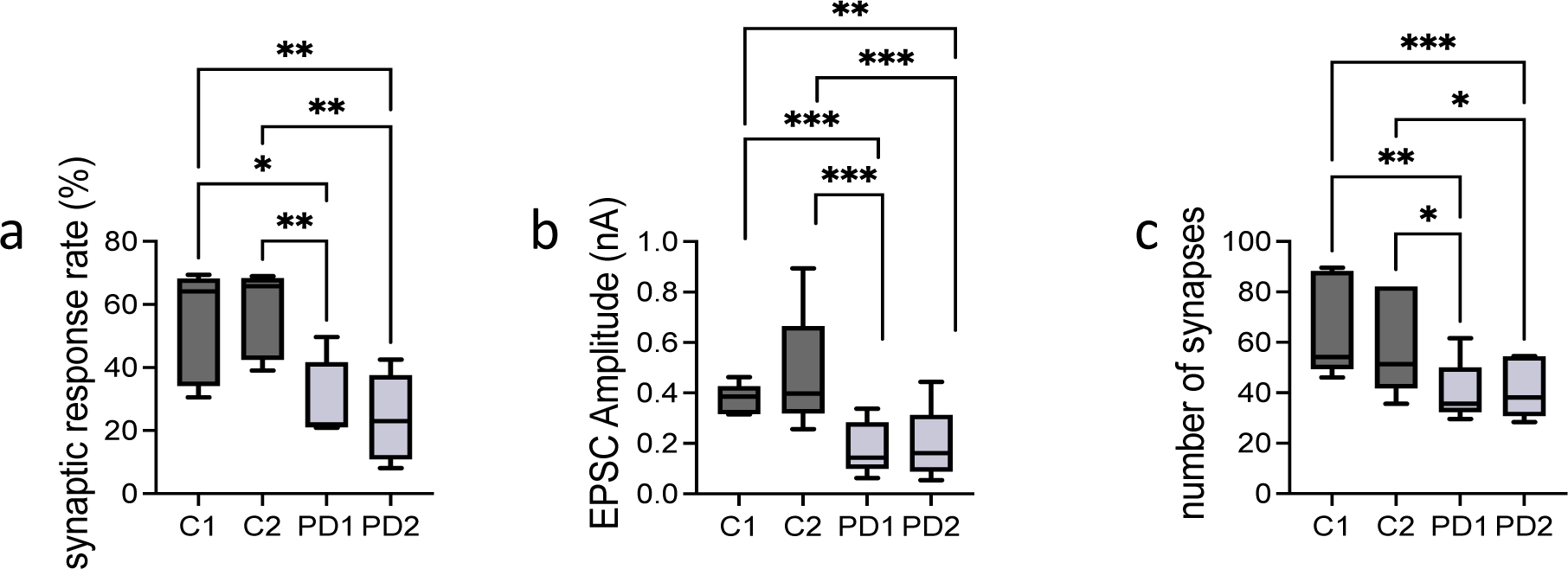
Synaptic response and number of synapses. Subgroup comparison of the proportion of neurons with AP-induced synaptic release, EPSC amplitude and number of synapses; (**a**) Synaptic response rate (‘subgroup’ factor: *p* = 0.0002, C1 vs PD1: *p*= 0.0229, C1 vs PD2: *p* = 0.0039, C2 vs PD1: *p* = 0.0071, C2 vs PD2: *p* = 0.0011, 2-way ANOVA followed by multiple comparison test Tukey), (**b**) EPSC amplitude (‘subgroup’ factor: *p* =<0.0001, C1 vs PD1: *p* = 0.0005, C1 vs PD2: *p* = 0.0013, C2 vs PD1: *p* = 0.0004, C2 vs PD2: *p* = 0.001, 2-way ANOVA with multiple comparison test Tukey). (**c**) Number of synapses (‘subgroup’ factor: *p* = <0.0001, C1 vs PD1: *p* = 0.0011, C1 vs PD2: *p* = 0.0009, C2 vs PD1: *p* = 0.0243, C2 vs PD2: *p* = 0.0264, 2-way ANOVA with multiple comparison test Tukey).

**Supplementary Figure 6.**
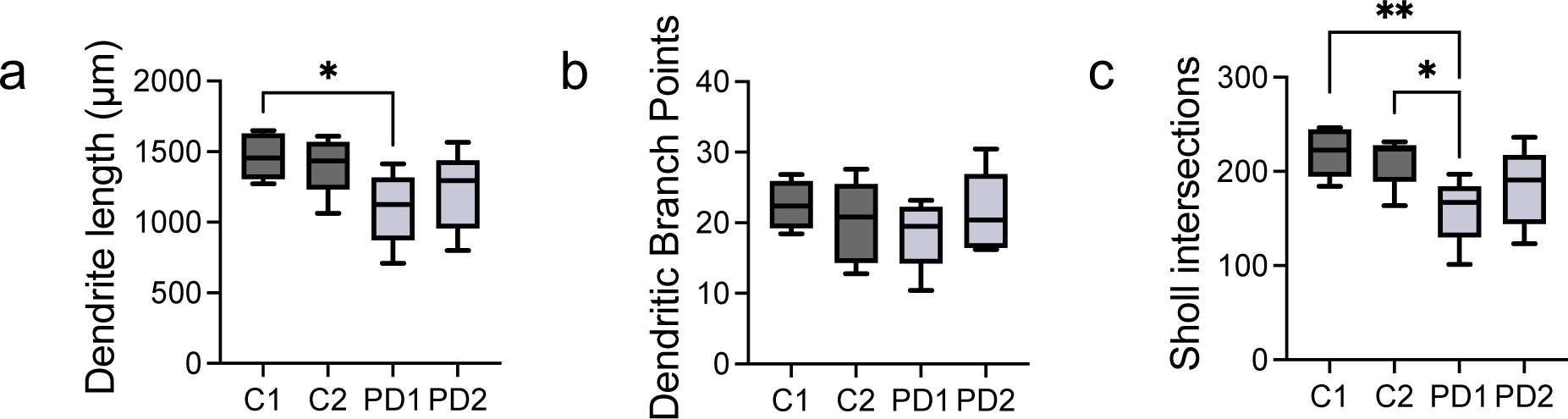
Morphological Analysis with immunohistochemistry. Subgroup comparison for summed up dendrite length, number of dendritic branch points and Sholl intersections. Both PD patient cell-lines had shorter overall dendrite length (**a**) and average numbers of dendrites intersecting Sholl circles up to 200 μm distance from the soma (**c**) (‘subgroup’ factor: *p* = 0.0119, C1 vs PD1: *p* = 0.0192, **c**, subgroup’ factor: *p* = 0.0051, C1 vs PD1: *p* = 0.0073, C2 vs PD1: *p* = 0.0355, 2-way ANOVA with multiple comparison test Tukey. Number of branchpoints did not differ clearly between subgroups (**b**).

